# Hybrid Generative Model: Bridging Machine Learning and Biophysics to Expand RNA Functional Diversity

**DOI:** 10.1101/2025.01.20.633900

**Authors:** Vaitea Opuu

## Abstract

Functional RNAs perform diverse catalytic roles, yet natural sequences represent only a narrow subset of what is possible. Rediscovering such activities requires exploring functional sequence diversity beyond natural RNAs. We introduce a hybrid generative model that combines a coevolutionary likelihood with an RNA secondary structure prior. This approach disentangles folding constraints from functional signals, enabling targeted diversification. On synthetic benchmarks, the model generates functional sequences beyond the training distribution. On large-scale ribozyme data, it improves the detection of active sequences and enhances sensitivity to local tertiary contacts. Finally, we introduce structural imprinting, a sampling strategy that uses alternative secondary structures to steer generation across under-sampled regions of sequence space. These results show that folding-informed generative modeling improves RNA design by supporting both extrapolation and control.

## Introduction

RNA plays essential roles in cellular processes, acting as both an information carrier and a biocatalyst in translation, splicing, and gene regulation [5]. These functions support the RNA world hypothesis [3], which also posits that RNA once carried out catalytic activities now performed by proteins. This motivates exploration of RNA sequence space beyond what is observed in nature [4], with applications in biosensing [6], gene modulation [23], and RNA-based therapeutics [22].

RNA design requires generating sequences that satisfy both structural and functional constraints. However, the size of sequence space and the scarcity of functional data make this difficult. Physics-based models offer generalizability but are limited by simplifications [11]. In contrast, ML models trained on MSAs can capture evolutionary patterns but tend to overfit and fail to extrapolate beyond the training distribution. Recent work has shown that generalization performance scales with sequence similarity to training data, underscoring the difficulty of cross-family modeling and supporting the utility of family-specific models for RNA design [18]. Moreover, family-specific models have demonstrated high accuracy in both protein and RNA design settings [19, 1, 7], motivating our focus on a single-family modeling strategy in this work.

The folding of RNA is largely governed by base-pairing, which allows efficient prediction of 2D structures from sequence [8]. This tractability has enabled hybrid models that combine ML with structural information. Prior work has used folding stability to reweight the sequence design a posteriori [7], improving the design of functional variants. However, these models do not incorporate folding energetics during learning, and typically assess success through sequence-level metrics. As a result, the structural diversity of generated sequences—i.e., the ability to explore alternative folds—remains largely unaddressed.

A common hypothesis in RNA design is that a functional RNA must be thermodynamically stable in its active 2D conformation to achieve function. This motivates integrating folding energetics directly into generative models. First, this can improve prediction of functional molecules by discriminating sequences that match evolutionary statistics but fail to adopt the correct fold. Second, folding energy depends only on the sequence and structure, not on the MSA or functional annotations. As such, it provides an external signal that can improve generalization, particularly when sampling away from the training distribution.

Similar strategies have proven effective in protein modeling. In particular, combining physical priors with evolutionary models has improved protein complex prediction [20], domain redesign [16], and enzyme reengineering [15]. These results suggest that integrating structural constraints into generative models can enhance both foldability and function.

Here we introduce a hybrid generative model with two contributions. First, the 2D folding energy is integrated directly into the training of a covariation model (or Potts model), allowing joint learning of evolutionary couplings and structural preferences. Unlike models that assume a structural configuration [21], we incorporate per-sequence folding energies as a biophysical prior. This reshapes the learned coupling landscape, increasing sensitivity to tertiary contacts. Second, we propose a new design paradigm based on structural diversification. By modulating the folding prior during sampling, we guide the model to generate sequences folding into alternative configurations not present during training. This enables extrapolation not only in sequence space but also in structure space.

We validate this approach on synthetic ligand-binding RNAs and on 8,000 Azoarcus ribozyme variants with measured activity [7]. The hybrid model improves ranking of active variants at high mutational distances. Unlike large neural networks, covariation models are interpretable and have long been used to infer structural contacts, providing an additional rationale for using simple, family-specific models in this context. Structural imprinting enables generation of stable sequences folding into distinct configurations, marking a shift from sequence-centered to structure-aware RNA design.

## Modeling

### Hybrid energy-based model

We define a generative energy-based model over RNA sequences that integrates evolutionary and thermodynamic constraints. The key assumption is that a sequence is functional only if it folds into a specific secondary structure *c*, which is fixed and assumed to support the target function. This constraint defines a biophysical prior over sequences based on their folding energetics. We then refine this prior using information from an alignment of functional homologs, resulting in a model that favors both stable and statistically consistent sequences. The probability of a sequence *S* is defined by the following energy-based model:

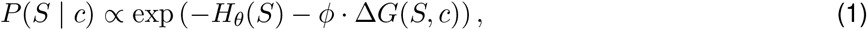

where *H*_*θ*_(*S*) is a statistical energy representing covariation and site preferences observed in the multiple sequence alignment (MSA), and Δ*G*(*S, c*) is the free energy of folding sequence *S* into structure *c*. The scalar parameter *ϕ* sets the relative importance of the structural stability constraint.

#### Statistical energy

The statistical energy *H*_*θ*_(*S*) is given by a Potts model over the sequence:

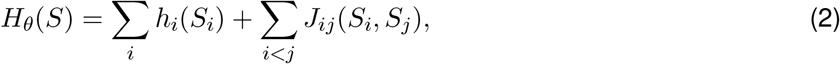

where *h*_*i*_ are position-specific fields and *J*_*ij*_ are pairwise coupling terms. These parameters define the distribution of sequences consistent with the observed covariation patterns in the MSA. They are not trained in isolation but are optimized jointly with the structural term as part of the full model.

#### Structural energy

The folding energy Δ*G*(*S, c*) is computed using the nearest neighbor thermodynamic RNA model [10], representing the predicted stability of sequence *S* when constrained to fold into the fixed structure *c*. This term encourages the model to assign higher probability to sequences that are thermodynamically stable in the functionally relevant conformation *c*.

#### Learning and sampling

All parameters, including *θ*= {*h*_*i*_, *J*_*ij*_} and *ϕ*, are optimized jointly by maximizing the likelihood of observed functional sequences under the full model distribution. The gradient of the loglikelihood with respect to the structural weight *ϕ* is given by:

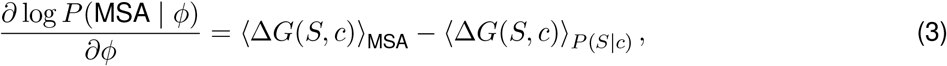

where the first expectation is over the empirical sequences in the MSA, and the second is estimated from the model distribution using MCMC.

This framework enables flexible generation and evaluation of sequences that simultaneously satisfy statistical and structural constraints.

### Illustrative validation on generated data

We first validate the hybrid model on artificial data with controlled structure and function to test if it performs as expected. This setup allows us to assess whether the model can disentangle folding stability from functional constraints, a key challenge in RNA design.

We generated an artificial dataset based on a 29-nt structured RNA designed to bind a four-nucleotide ligand (configuration A, Fig. 2). This short construct was chosen to enable comprehensive sampling and evaluation. Random in silico mutagenesis was followed by selection for both folding stability and binding affinity, computed with ViennaRNA. We retained *<* 400 sequences satisfying predefined folding and binding thresholds to form the training set. To account for the narrow mutational range of this data, we introduced a regularization term *ω* (see SI), which penalizes divergence from the training distribution under the assumption that mutations reduce function exponentially. Similarly to the relative temperature of the structural prior *ϕ, ω* is a tunable parameter optimized during training. Binding scores were computed based on predicted ligand interaction energy at the 5 end, using McCaskill algorithm (details in SI). Fig. SI2 shows that HYB outperforms REG in generating active sequences across mutational distances.

**Fig. 1:**
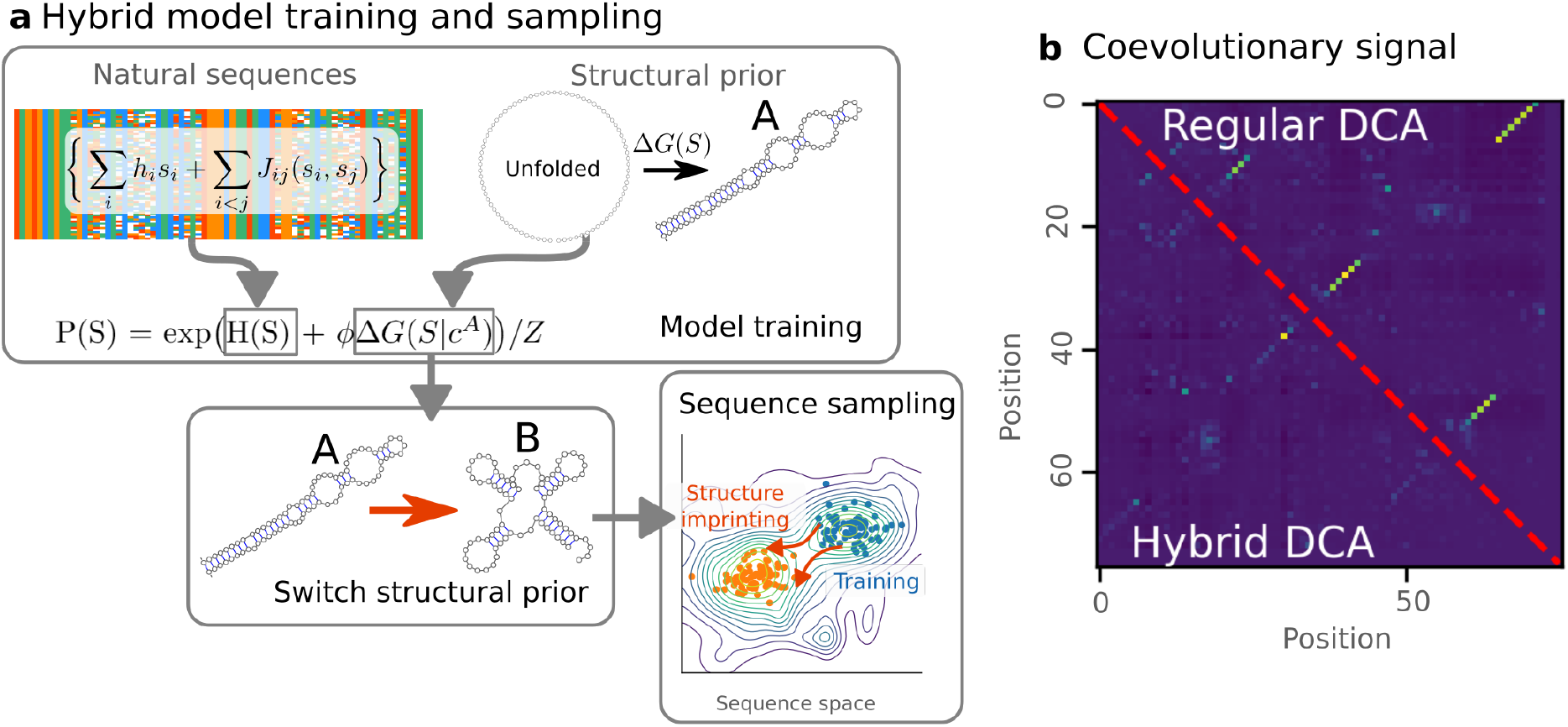
Hybrid DCA model integrating structural priors. (a) Schematic overview of hybrid DCA. Natural sequence alignments inform DCA parameter estimation (top left), while a structural prior (top right) encodes 2D folding stability. The decoupled formulation allows alternative priors to introduce sequence diversity (bottom left), enabling sampling of sequences folding into different structures while retaining statistical constraints from the MSA (bottom right). (b) DCA coupling matrices for the tRNA family (RFAM): regular DCA (top triangle) and hybrid DCA (bottom triangle). In hybrid DCA, 2D signal is reduced due to its explicit incorporation in the prior.

**Fig. 2:**
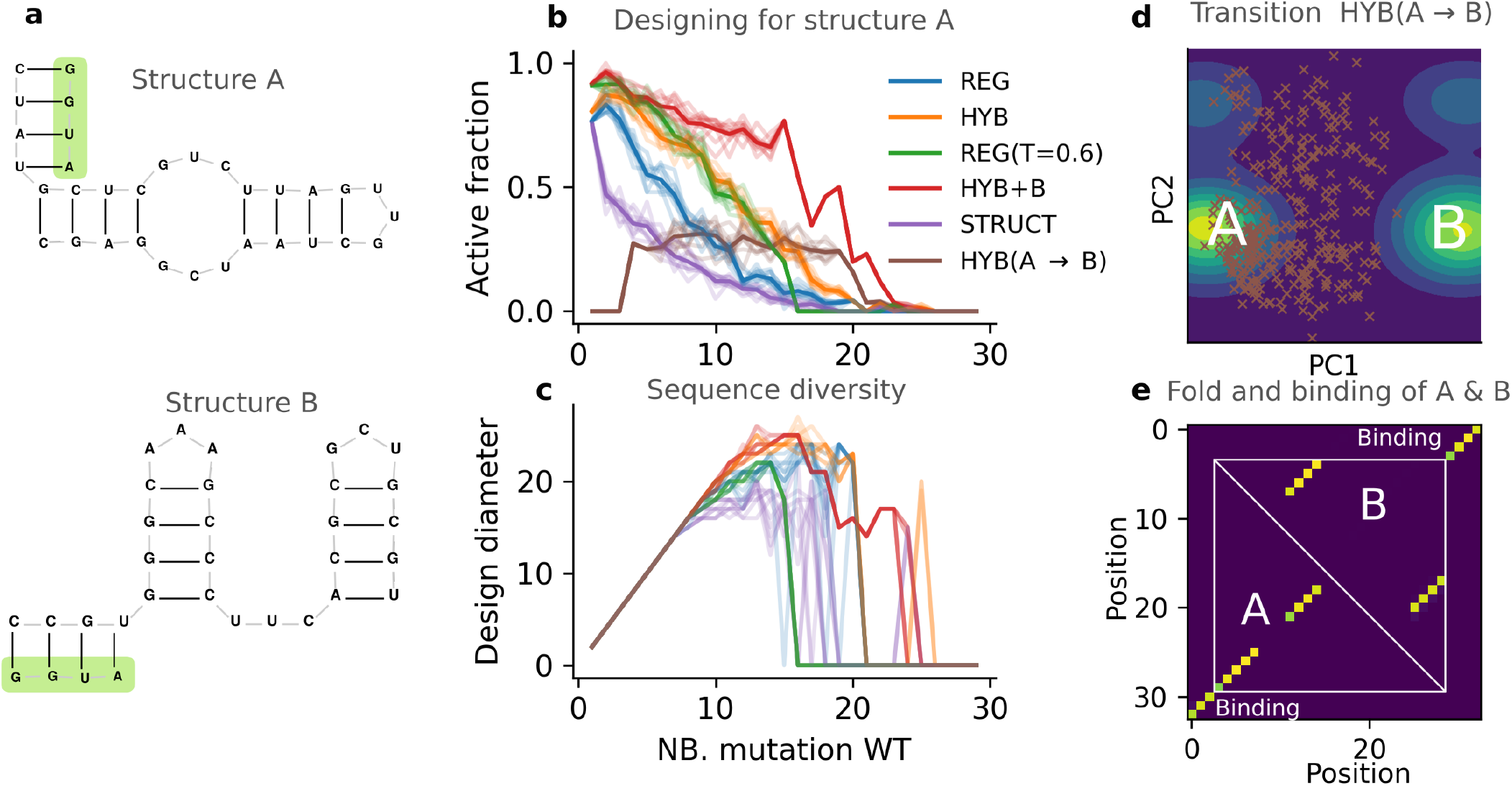
Disentanglement of stability from function on artificial data. (a) Secondary structures of configurations A and B used in design. The ligand is shown in green. (b) Success rate of design methods. The plot shows the fraction of active sequences as a function of their number of mutations to the artificial wild type (WT). For each mutation distance, we randomly selected 100 sequences designed by each method (REG = regular DCA; HYB = hybrid model using configuration A; REG(T=0.6) = regular DCA at lower sampling temperature; HYB+B = hybrid model with increased data-driven weight; HYB(A→B) = hybrid model using configuration B). The process was repeated 10 times to estimate variability. (c) Sequence diversity. For each mutation distance, we computed the number of mutations between the most dissimilar designed sequences (design diameter). (d) Interpolation between binders in configuration A and B. PCA was performed on training sequences (density shown as contours; brighter means higher density). HYB(A→B) designs are shown as brown crosses. (e) Base pair probability matrices of successful designs for both configurations.

Although the training data contains sequences with at most ten mutations from the wild type, HYB generates functional binders up to twenty mutations away, outperforming the regular DCA model (REG), which lacks structural information (Fig.2). At 15 mutations, HYB achieves 22.5% successful designs versus 8% with REG, indicating its ability to extrapolate beyond the training distribution. A potential explanation for this improvement is reduced diversity, which can increase activity by collapsing designs around high-fitness peaks. To test this, we computed the design diameter—the maximum Hamming distance between any two designed sequences at fixed mutational depth. As shown in Fig.2c, HYB maintains diversity comparable to REG. As expected, lowering the sampling temperature in REG (REG(*T* = 0.6)) increases the binder fraction but collapses sequence diversity. In contrast, HYB achieves high activity without sacrificing diversity. To dissect the respective roles of folding and binding, we varied the weight of the data-driven term in HYB. Increasing this weight improves the binder fraction without compromising diversity (Fig. 2). In contrast, STRUCT, a baseline sampling only from the folding prior, produces stable structures but few binders, confirming that folding energy alone does not encode binding specificity. These results show that HYB enables controlled generation of diverse, high-activity sequences beyond the training space.

Finally, we asked whether the functional constraints captured by HYB can transfer to new and structurally distinct contexts. We isolated the data-driven component from a HYB model trained on configuration A and combined it with a folding prior for a distinct conformation B, which shares no base pairs with A and was not seen during training. This strategy steers sampling toward B while preserving the binding signal from A. As shown in Fig.2e, the resulting sequences retain binding activity and fold into B. Folding into B was confirmed using the McCaskill algorithm. To visualize how the sequences relate to those of A and B, we projected all sets in a PCA space built from one-hot encoded sequences sampled from each model. Fig.2d shows that these sequences lie between the A and B binders in this projection, illustrating a continuous and controlled shift. While this is a conceptual visualization, it suggests that functional imprinting is feasible and tunable when model components are modular.

Together, these results show that HYB enables functional optimization independently of structural stability, supporting both controlled diversification and extrapolation in RNA design. The next step is to evaluate whether these capabilities hold in real experimental settings, where functional constraints and structural variability are less controlled.

### Improved detection of ribozyme functionality

A central challenge in RNA design is to generate diverse sequences that preserve function. We hypothesize that hybrid models—combining ML with explicit folding priors—enhance functional sequence detection by increasing sensitivity to structural constraints, including local tertiary interactions.

To evaluate our hypothesis, we used a large-scale experimental dataset from [7], where over 30,000 Azoarcus ribozyme variants were generated using different generative strategies and tested for catalytic activity. We focused on three strategies spanning statistical to structurally informed approaches: regular DCA trained on the same MSA (Data-DCA); a naive hybrid model combining DCA with a 2D structural prior (Data-HYB); and a base-pair replacement model incorporating fixed tertiary contacts from the 1U6B structure (Data-BPR3D) [**?**]. Sequences were labeled functional or non-functional using the activity threshold defined in the original work, yielding 33% functional sequences among 8,097 variants. The dataset spans mutational distances from 0 to over 100 substitutions relative to the wild type.

We next assessed whether the models can identify functional sequences by evaluating their ability to rank active variants higher than inactive ones. DCA models were trained on the provided MSA in [7] and used to assign statistical energies, which served as classifier scores. For the hybrid models (HYB), which combine DCA with either the consensus 2D structure (ALI) or the experimental structure (EXP), we computed the classifier score as the sum of DCA and folding energies. Folding energy alone (STA) was used as a baseline score, assigning lower scores to sequences with low free energy, thereby favoring RNAs predicted to stably fold into the target structure. Classification refers here to ranking sequences by model-assigned scores and comparing them against binary functional labels. Performance is assessed using average precision. Statistical significance was assessed using a one-sided Student’s t-test (*α* = 0.05) on 30 bootstrap samples per bin, testing the hypothesis that HYB outperforms the structure-naive DCA (REG). All models are evaluated in an unsupervised setting unless otherwise noted. Supervised classifiers are introduced below for context, not comparison.

We now turn to the results to test whether hybrid models improve the identification of functional sequences compared to structure-naive baselines. First, we tested the classification in Data-DCA but no systematic gain in performance was observed. These sequences are sampled based on the evolutionary couplings learned from the MSA, biasing them toward regions close to the training distribution. As a result, HYB and REG perform similarly in most bins, with modest gains only beyond 50 mutations. In contrast, sequences from Data-HYB deviate more from the training set. In this case, HYB outperforms REG in more than half of the bins above 30 mutations, as indicated by stars in Fig. 3a marking statistically significant differences. This supports the hypothesis that the structural prior enables better generalization to distant sequence space. This trend is further amplified in Data-BPR3D, where mutations are introduced randomly but restricted to positions compatible with the 2D structure, while nucleotides involved in tertiary contacts are fixed to the wild-type. Here, the added structural constraints preserve functional signal in more divergent backgrounds. The synergy between the data-driven component and structural prior is further illustrated by the poor performance of the STA score, which is systematically lower than both HYB and REG across all bins, highlighting the insufficiency of folding energy alone in capturing functional specificity. To contextualize these results, we compared with a 2-layer perceptron trained on labeled data across mutation bins (SI Fig. 5). While this supervised model outperforms all unsupervised baselines, it relies on labeled activity data and is not directly comparable to the DCA-based generative models.

**Fig. 3:**
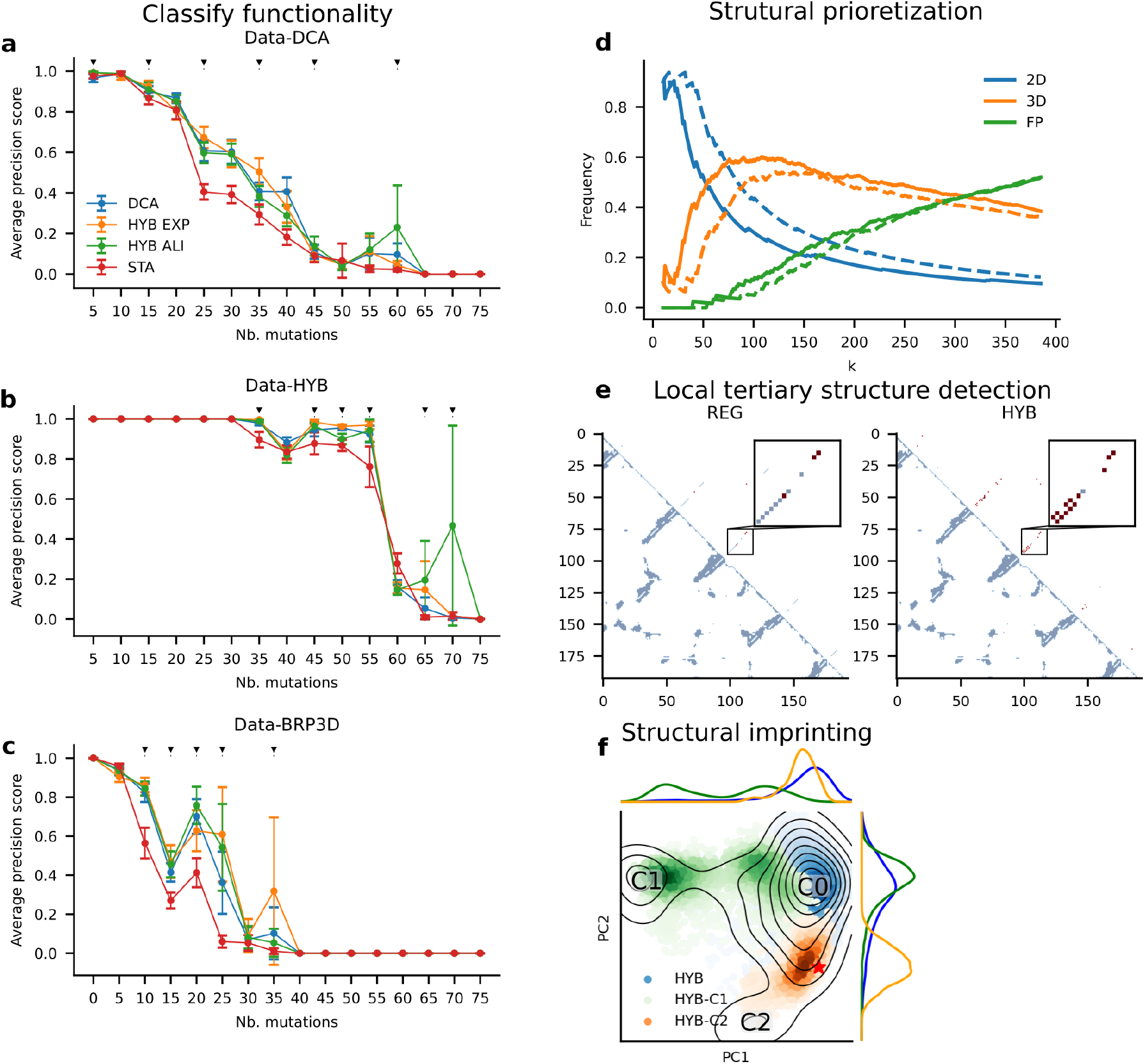
Hybrid models improve functional classification, structural interpretability, and enable targeted exploration of undersampled sequence space. (a–c) Average precision (AP) scores for classifying functionality across mutation bins (5-mutation intervals) for three datasets: Data-DCA (a), Data-HYB (b), and Data-BPR3D (c). HYB variants outperform DCA in several bins (black triangles, p¡0.05, one-sided t-test, bootstrap n=30). (d) Structural contact enrichment as a function of top-ranked DCA couplings (k). HYB improves tertiary contact (3D) prioritization at k=118, while DCA better recovers 2D base pairs at k=40. (e) Contact maps show coupling signal from DCA (left) and HYB (right), with HYB redistributing signal to short-range tertiary contacts (red). Insets highlight tertiary motifs. (f) PCA of sampled sequences. HYB designs populate underrepresented clusters (C1, C2) not covered by DCA alone. Clusters were manually defined from the natural MSA projection.

To better understand why the hybrid model improves classification performance, we analyzed the structural features it captures, focusing on tertiary contacts, which are critical for catalysis. We hypothesize that the hybrid model improves activity detection by shifting focus toward tertiary contacts. We define tertiary contacts as nucleotide pairs with at least one heavy-atom distance below 12 Å, a cutoff commonly used in force fields to neglect electrostatic interactions beyond this range. Known 2D contacts identified by DSSR were excluded from the tertiary contact set. To infer contacts, we compute pairing scores using the Frobenius norm of the coupling terms in the DCA parameters. We rank all pairs by score and retain the top *k*. Enrichment is defined as the proportion of true tertiary contacts recovered within the top-k ranked pairs. In Fig. 3d, we varied *k* and observed that top-scoring pairs are more enriched in tertiary contacts in HYB than in REG, with peak enrichment at *k* = 118 for HYB and *k* = 140 for REG. Concurrently, detection of base-pair (2D) contacts is lower in HYB compared to REG, with the largest difference at *k* = 40, consistent with a redistribution of coevolutionary signal from secondary to tertiary interactions. The HYB-inferred contact map shows detection of local but not long-range tertiary interactions (Fig. 3), suggesting that structural priors steer the model toward capturing short-range 3D constraints. These results indicate that the hybrid model attenuates 2D signals while enhancing sensitivity to 3D interactions, that we investigate further in the next section. Regardless of direct impact on classification, the shift toward tertiary contacts highlights the structural interpretability gained by integrating folding priors.

Having established improved classification performance, we now show how the hybrid model can be used to increase diversification of generated sequences, with a focus on diversity and coverage in sequence space. Here, we use the HYB model trained with the ALI 2D configuration prior. To visualize the diversity of sampled sequences, we projected them onto the principal components of the MSA sequence features, computed with PCA on one-hot encoded sequences. We define “diversity” as the spread of generated sequences across sequence space, and “coverage” as the proportion of generated sequences occupying predefined clusters (C0, C1, C2) in the principal component projection. Figure 3 shows that DCA primarily samples the most populated cluster (C0) from the natural MSA sequences, while the other two less populated clusters (denoted C1 and C2) remain sparsely sampled, as expected. For clarity, clusters C0, C1, and C2 were defined by manually applying thresholds on the first two principal components to separate visually distinct groups in the projection. To improve coverage, we leveraged the hybrid model’s separation of structure and data-driven energy to bias the sampling toward sequences predicted to adopt the 2D structure characteristic of C1, while maintaining low hybrid energy. To define the 2D configurations associated with C1 and C2, we used RNAalifold on sequences sampled in each cluster. We then applied the structural imprinting strategy described above, replacing the folding energy term for the global ALI configuration with the folding energy into the C1 configuration, weighted by a factor *ϕ* = 2 to enforce structural bias. The resulting set (HYB-C1 in Figure 3f) now spans the region corresponding to C1, increasing cluster occupancy from 4% with the global ALI prior to 43% with the C1 structural prior. We applied the same approach to C2, demonstrating the generality of the strategy. Figure 3f shows that HYB-C2 also fills the C2 region sparsely populated by DCA, increasing occupancy from 1.3% to 13%. In both cases, the hybrid model increased coverage of underrepresented regions in sequence space relative to DCA alone.

We propose a hybrid modeling strategy to enhance the diversity of functional RNAs. Re-analysis of experimental data shows that incorporating 2D folding priors improves the identification of active sequences, particularly at high mutational distances. This improvement correlates with increased detection of local tertiary interactions. Finally, we show that tuning the structural prior enables targeted sampling of underexplored regions, providing a flexible approach to diversify RNA designs.

### Assessing Structural Priors in Contact Inference

DCA often recovers RNA 2D contacts from MSAs, partly because conserved base pairs influence alignment construction, e.g., via covariance models used by INFERNAL [13]. In contrast, tertiary (3D) contacts, which exhibit weaker coevolution, remain challenging to detect.

We evaluate whether the HYB approach improves tertiary contact inference. For this, we used the curated dataset of RFAM families of homologous natural RNAs described in [17]. Following [24], we selected 57 families, with sequence lengths ranging from 41 to 496 nucleotides. Each RFAM family is associated with a corresponding 3D structure from the PDB, which we used to derive the experimental 2D structure via DSSR [9]. The MSAs were trimmed to retain only positions aligned with the resolved 3D structure, defining the target sequence. In the 3D structure, two nucleotides were considered in contact if any of their heavy atoms were within 12 Å. We defined 3D contacts as those not present in the 2D structure, including pseudoknots.

We trained one HYB model and one regular DCA model (with *ϕ* = 0) for each of the 57 selected families. For the structural stability term in the HYB model, we considered three structure sources: EXP (the experimental 2D structure), MFE (the minimum free energy structure predicted for the target sequence), and ALI (the consensus structure from the MSA via RNAalifold). For each training, the correlation between empirical and model pairwise nucleotide frequencies exceeded 0.8. We enforced 2D priors by assigning maximum coupling scores to 2D contacts, ensuring their inclusion in the predicted contact set. We then ranked all position pairs by coupling strength and selected the top contacts across varying fractions of the sequence length *L*. Predic-tion performance was measured using the positive predictive value (PPV), defined as the fraction of predicted contacts that match the true 3D contacts.

We evaluated contact prediction across all types using the hybrid model (HYB) and the regular DCA model (REG, with *ϕ* = 0) over 57 RNA families. HYB yields higher PPVs than REG when using the ALI prior, across contact fractions from 0.1 to 2 *L* (Fig.4; Wilcoxon test, *p <* 0.05). REG shows a gradual decline in PPV from 0.82 to 0.39 as more contacts are considered, suggesting limited capacity to capture weaker signals beyond the most conserved contacts. The differences observed with HYB reflect the degree to which structural priors align with the sequence distribution in the MSA.

**Fig. 4:**
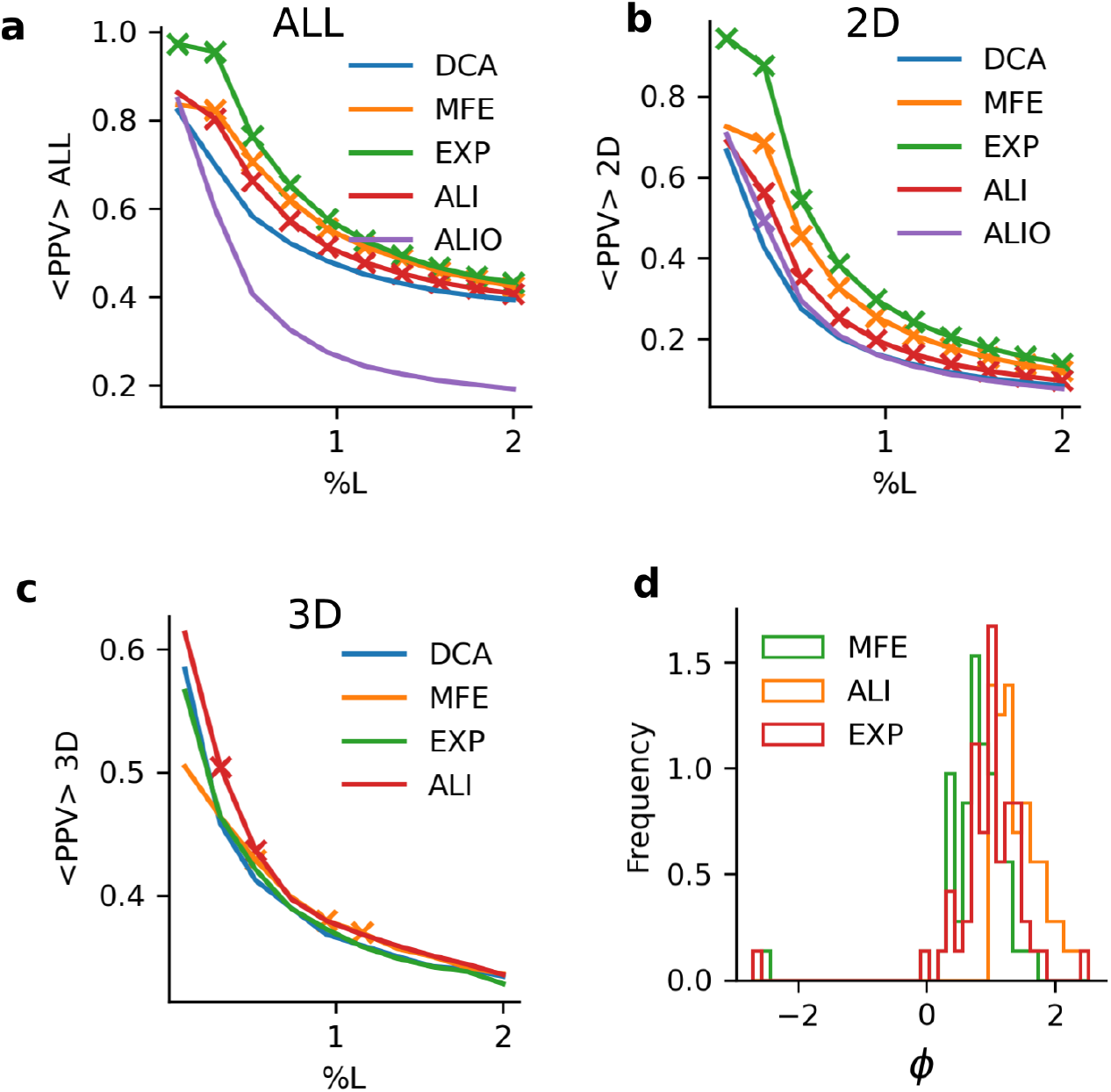
Improved detection of tertiary contacts with hybrid models. (a–c) Contact prediction accuracy measured by PPV as a function of top-ranked contact pairs (expressed as %L, sequence length). ALL (a), 2D base pairs (b), and 3D tertiary contacts (c) are shown for regular DCA (blue) and hybrid models using MFE (orange), EXP (green), ALI (red), and ALIO (purple) structural priors. Crosses mark %L values where a hybrid model significantly outperforms DCA (Wilcoxon test, p ¡ 0.05). (d) Distribution of contact enrichment scores () across all tested families for hybrid models using MFE, ALI, and EXP structural priors.

In the case of 2D contacts, REG reaches PPVs between 0.66 and 0.08, comparable to RNAalifold (0.70 to 0.07), confirming that 2D contacts are well recovered from the coevolutionary signal. HYB yields higher values with the ALI and MFE priors, consistent with structural information contributing when aligned with conserved features in the MSA. The EXP configuration, which includes the known target structure, defines an upper bound and is excluded from comparison. Since most 2D contacts are already captured by folding models such as RNAalifold, only limited improvement is expected from hybrid modeling in this regime.

Among all contact types, 3D contacts remain the most difficult to infer due to weaker coevolutionary signals in MSAs. REG achieves PPVs ranging from 0.58 to 0.28, comparable to those of HYB. With the ALI prior, HYB recovers a slightly greater fraction of 3D contacts (PPVs from 0.61 to 0.33), with significant but marginal improvements at %*L* = 0.31 and %*L* = 0.5. The MFE prior also yields significant gains at %*L* ∈ {0.31, 0.94, 1.1}. Unexpectedly, the EXP prior, which contains the true target structure, does not improve 3D prediction. A possible explanation is that EXP diverges from the MFE configuration and does not align with the free energy landscape encoded in the MSA. The influence of the structural prior is captured by the parameter *ϕ*, which is consistently higher when the prior aligns with the MSA and lower otherwise (ALI aligns significantly better; t-test, *p <* 10^*−*5^; Fig.4d). This supports the idea that structural priors consistent with the alignment guide the model toward contacts not captured by coevolution alone. Non-generative variants such as mfDCA and PLM were also tested, but did not benefit from structural reweighting (see SI).

The hybrid model improves 3D contact prediction marginally, with gains depending on how well the structural prior aligns with both the MSA and the underlying energy landscape. By incorporating consistent structural information, it increases the weight on 3D interactions without altering overall prediction trends. This shift may ultimately support functional design, as 3D structure is often critical for catalysis.

### Structural Imprinting for Sequence Diversification

Having shown that the hybrid model improves classification and enriches for tertiary contacts, we now test whether structural priors can guide extrapolation beyond the training data, by targeting distinct sequence clusters within natural RNA families.

To test this, we selected 25 MSAs of natural RNA families from [17], each containing between 200 and 5000 sequences and with lengths below 300 nucleotides, as longer sequences are associated with reduced modeling accuracy [14]. We then clustered the sequences within each MSA into two groups using spectral clustering based on Hamming distance, as implemented in Scikit-learn (version 1.0.2). While the clustering is not guaranteed to produce functionally distinct groups, splitting based on sequence divergence tends to separate conserved motifs, which makes generalizing from one cluster to another non-trivial. For each cluster pair, we computed a consensus structure using RNAalifold, referred to as configuration A and B. The base-pair distance, defined as the number of base pairs present in one structure and not the other, was on average 7, indicating that most MSAs contain structurally consistent but non-identical sequences. Our goal is to assess whether a model trained on one sub-MSA can generate sequences representative of the other. Success in this task would suggest that structural variation can guide extrapolation toward unseen but functionally relevant RNAs.

To establish a baseline, we trained one regular DCA model (REG) per cluster, resulting in 25 × 2 instances. From each model, we generated 10^6^ MCMC samples, initialized with the cluster consensus sequence, and randomly selected 500 sequences for analysis. To quantify extrapolation, we projected all sequences onto a PCA space constructed from the one-hot encoded natural sequences of each family, retaining principal components that capture 90% of the variance. This projection preserves the major sequence variation across the family and provides a low-dimensional representation to compare generated and natural sequences. We then estimated an empirical density for each cluster using kernel density estimation (KDE) implemented in Scikit-learn, with a Gaussian kernel and a bandwidth parameter set with the scott method. For each generated sequence, we computed a p-value under the KDE of the opposite cluster using Monte Carlo integration, sampling 10^4^ points from the KDE distribution to approximate the integral. Sequences with p-value *>* 0.01 were considered successful cross-samples, indicating that the generated sequence falls in a high-density region of the opposite cluster and is statistically consistent with its sequence distribution.

Regular DCA sampling reveals limited extrapolation between clusters. Our results showed that the regular model (REG) generated at least one successful cross-sampled sequence in 14 out of 50 cluster pairs, for a total of 353 successful cross designs (Fig. 5d). This indicates that extrapolation beyond the training cluster is possible but limited. These cases correspond to clusters that are not fully isolated in sequence space, suggesting the presence of mutational paths between them. Supporting this, the average base-pair distance between their consensus structures is approximately 3, compared to 7 on average across all cluster pairs. This reduction confirms that successful extrapolation by REG requires low structural divergence between clusters. Increasing the sampling temperature (*T*) reduced the number of successful designs to 160, and applying a bias term (*ω*) lowered it to 217, confirming that naive diversification strategies without guidance are ineffective. We next tested whether structural guidance enables broader extrapolation. To evaluate this, we applied the hybrid model (HYB), trained on one cluster using its consensus structure, and sampled sequences using the folding energy of the opposite cluster. This “structural imprinting” aims to steer the model toward generating sequences representative of unobserved clusters. For each family, we trained HYB on both clusters, extracted the data-driven component, and sampled sequences with structural priors from the opposite cluster using *ϕ* = 2 and *ϕ* = 4. From each, we randomly selected 500 sequences. Across all families, HYB generated 624 successful cross-samples in 15 cluster pairs (Fig.5d), improving both the number and diversity of successful extrapolations. Importantly, HYB consistently recovered cross designs in the same families as REG but yielded more successes per pair, suggesting that structural priors enhance sampling across functional modes. The difference in yield is statistically significant (Wilcoxon test, p=0.03). Representative cross-sampled sequences are shown in Fig. 5a,b.

**Fig. 5:**
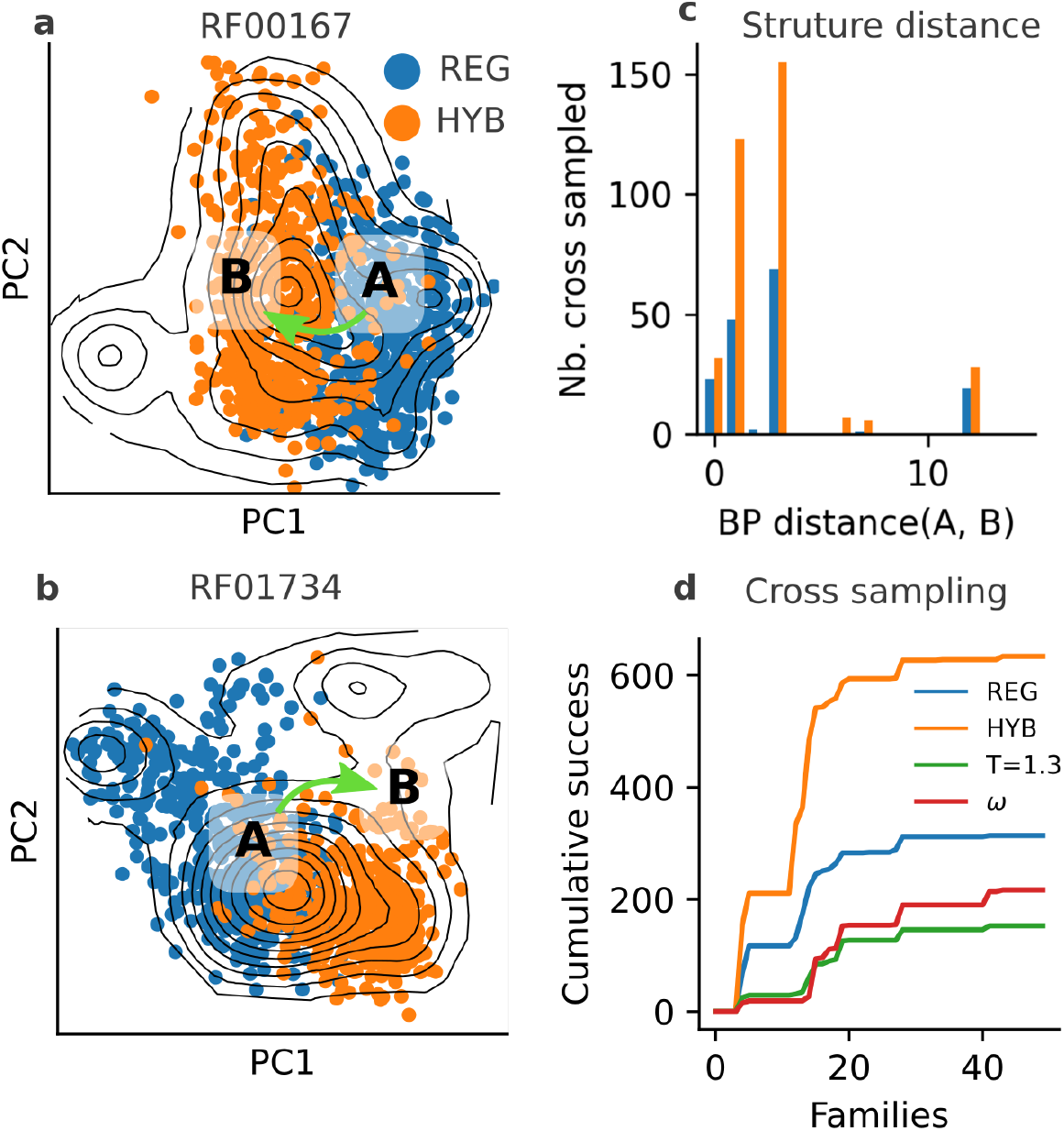
Generalization to unseen sequences via structure imprinting. (a, b) PCA projections of natural sequences in two RNA families (RF00167 and RF01724), with density contours indicating sequence distributions. Cluster centers A and B are annotated. REG (blue) designs reproduce the training cluster, while HYB (orange) samples into the unseen cluster using the alternative structural prior. Green arrows indicate extrapolation direction. (c) Number of successfully cross-designed sequences as a function of base-pair distance between configurations A and B, showing HYB generalizes better across increasing structural differences. (d) Cumulative count of successful cross-sampled sequences across 45 RNA families under a kernel density p-value threshold of 0.01. HYB outperforms REG and other baselines (T=1.3, *ω*-scaling) in sampling into unseen clusters.

While HYB doubled the success rate, it is not without limitations. The first caveat is that HYB still requires some overlap between the training and target clusters, as seen above. This is because successful extrapolation occurred mostly in cluster pairs where REG already achieved partial success, indicating shared motifs or mutational interfaces. The second limitation concerns the structure: HYB performs best when the consensus structures of the two clusters differ modestly (3.4 base pairs on average, Fig. 5c). Identical consensus structures offer no new directional signal and merely reproduce the training distribution, as observed for REG. Conversely, when the structures differ too strongly (more than 10 base pairs), the sampling fails to produce sequences statistically consistent with the opposite cluster. These results suggest that successful cross-sampling depends on the presence of a smooth structural gradient: too little signal provides no guidance, too much acts as a disruptive force, destabilizing the sampling process.

Generating functional sequences beyond the training set remains a central challenge in machine learning. We showed that the hybrid model enhances sequence generation across distinct subpopulations by leveraging small structural shifts. This suggests that structural guidance offers a viable path to uncover hidden functional diversity.

## Discussion

This work introduces a hybrid generative model that integrates evolutionary constraints learned from sequence alignments with an explicit folding prior, enabling controlled RNA design beyond observed data. By disentangling structural stability from functional covariation, the model addresses a central challenge in RNA modeling: the generation of diverse and functional sequences.

Across artificial and natural datasets, we show that the hybrid model improves detection of functional RNAs relative to structure-naive baselines. On synthetic ligands, it supports extrapolation far beyond the mutational scope of the training set without sacrificing diversity. On experimental ribozyme data, it improves classification performance at high mutational distances, where sequence divergence renders standard coevolutionary scores ineffective. These improvements reflect the capacity of the hybrid model to combine complementary sources of information—sequence statistics, structural constraints, and functional selection—into a single generative framework.

Mechanistically, the gains are explained by a redistribution of coevolutionary signal: when structural priors are introduced, the model reduces emphasis on canonical 2D contacts and increases sensitivity to tertiary interactions. This shift, confirmed across both experimental and curated structural datasets, suggests that folding constraints reweight the learning process toward local 3D dependencies that are otherwise difficult to access from MSA statistics alone.

Importantly, the hybrid formulation allows flexible control at generation time. Through structural imprinting, we steer sampling toward alternative RNA folds while retaining learned functional signals. This enables extrapolation across sequence clusters and conformations, which is not possible using DCA alone. However, imprinting remains effective only when structural differences are moderate—large shifts introduce constraints incompatible with the data-driven component, leading to sampling failure. This boundary represents how far hybrid models can generalize while preserving functional plausibility.

Overall, these results demonstrate that incorporating folding priors improves both inference and generation in data-scarce or divergent sequence regimes. Although the present implementation uses a standard 2D folding energy model, the framework is compatible with more detailed priors, such as kinetic models or approximations of atomistic force fields, as they become computationally tractable. Hybrid models therefore provide a practical approach for structure-informed RNA design, supporting interpretability and extrapolation in both evolutionary and synthetic contexts. More generally, the integration of mechanistic priors into generative models is likely applicable beyond RNA, including in large language models, where it may facilitate improved generalization or model compression by embedding structured constraints directly into the architecture.

## Code and data availability

The code and data used in this research is available at https://github.com/vaiteaopuu/rna design biophy ml. The training of the hybrid model finishes in seconds to minutes depending on the length of the MSA.

## Acknowledgments

I thank Philippe Nghe and Matteo Smerlak for helpful discussions. I thank Frederik Koch for doing the preliminary investigations.

## Supplementary Information

### Secondary Structure Inference with DCA

The DCA generative model was originally developed for protein contact prediction^∼^[12]. To infer contacts, we compute the Frobenius norm of the couplings at each pair (*i, j*) as

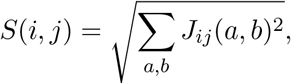

where *J*_*ij*_(*a, b*) is the pairwise coupling between nucleotides *a* and *b* at positions *i* and *j*, including the gap state. The top *k* residue pairs are selected based on APC-corrected scores^∼^[2], where *k* is set as a fixed multiple of the sequence length *L*, defined as the number of alignment columns after filtering. Performance is reported as a function of *k/L*.

### Biophysically Informed Mean-Field DCA

Mean-field DCA approximates the posterior over sequences by estimating coupling parameters from the covariance matrix:

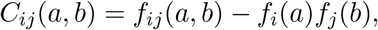

where *f*_*ij*_(*a, b*) is the joint frequency of nucleotides *a* and *b* at positions *i* and *j*, and *f*_*i*_(*a*), *f*_*j*_(*b*) are single-site frequencies. The couplings are inferred from the inverse of the covariance matrix:

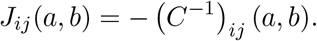

To reduce redundancy in the MSA, we apply sequence reweighting^∼^[12]. Additionally, to incorporate folding bias, we assume the empirical sequence distribution is influenced by the folding free energy Δ*G*(**S**, *c*). Since energy distributions vary across RNA families, we center and normalize Δ*G*(**S**, *c*) within each family, denoting the result as Δ*G*^*∗*^(**S**, *c*). The weight for each sequence **S** is computed as:

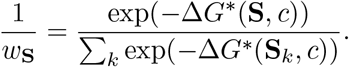

Frequencies are then computed using:

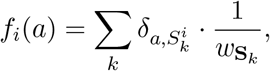

where *δ*_*a*_, 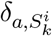 is the Kronecker symbol, equal to 1 if nucleotide *a* is present at position *i* in sequence *k*, and 0 otherwise. This variant is implemented using Python and NumPy (version 1.21.6).

### Bias Toward a Reference Sequence

Random mutations in functional RNAs typically cause exponential decay in activity. To model this, we introduce a sequence bias term *ω* that penalizes divergence from a wild-type (WT) sequence:

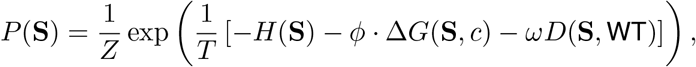

where *D*(**S**, WT) is the number of mutations relative to the WT. The parameter *ω* is updated via:

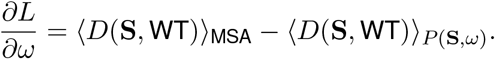

As with *ϕ*, the second term is approximated by the current sample at update time.

### Simulated Folding and Binding Experiment

We generated an artificial dataset to simulate binding and stability constraints starting from a functional RNA. A single-stem configuration, denoted A (see Fig.^∼^2), was designed to bind a short ligand (GGUA). The initial functional RNA sequence was optimized via MCMC to fold into configuration A and bind the ligand in a defined pose.

Sequence selection used the structure-based cost function from^∼^[7], which relies on the McCaskill algorithm to evaluate base-pairing probabilities within the receptor and with the ligand. The score penalizes alternative folds and alternative ligand poses, favoring sequences that are both structurally and functionally specific. The resulting wild-type (WT) RNA was then randomly mutated. For each mutation class (1–10 mutations), we generated 10,000 variants and retained those with a structure score *<* 0.6, yielding 342 mutants in total (Fig.^∼^SI^∼^4).

Binding was quantified as the probability that each position in the receptor’s binding site is correctly paired with the ligand nucleotides. This is also computed using the McCaskill algorithm. The resulting binding score reflects the ensemble-averaged specificity of the interaction. A sequence is considered active for binding if its score exceeds 3.

## Figure SI

**Fig. SI 1:**
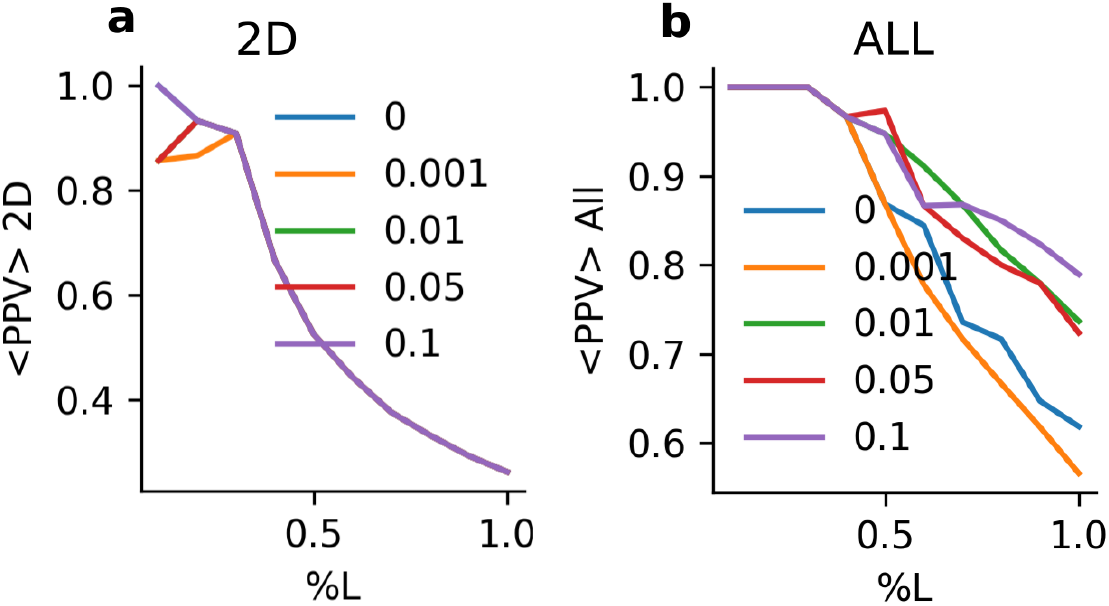
Effect of *L*_1_ regularization on DCA for the tRNA MSA family, shown for (a) 2D contacts and (b) all contacts.

**Fig. SI 2:**
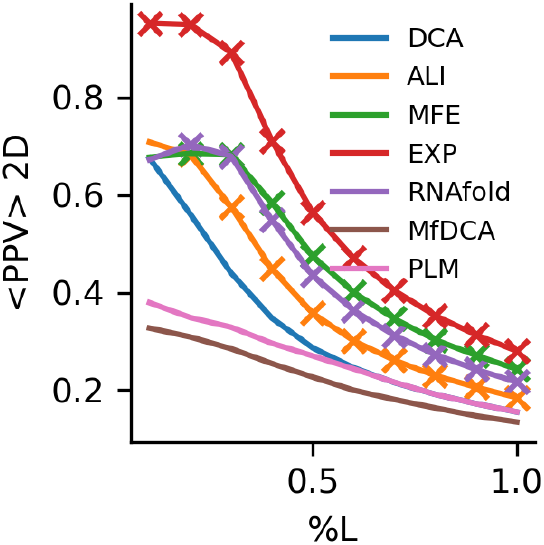
2D accuracy. Accuracy for 2D contact prediction as a function of ranked contacts based on scores *S*_*i,j*_. PPV is reported for regular DCA (blue), hybrid DCA with the ALI prior (ALI), MFE prior (MFE), EXP prior (EXP), RNAfold predictions (RNAfold), Mean Field DCA (MfDCA), and Pseudo-likelihood DCA (PLM). The x-axis shows the number of ranked contacts as a fraction of RNA length *L*; the y-axis shows PPV. Crosses mark cases where a method significantly outperforms regular DCA (Wilcoxon test, *p <* 0.05).

**Fig. SI 3:**
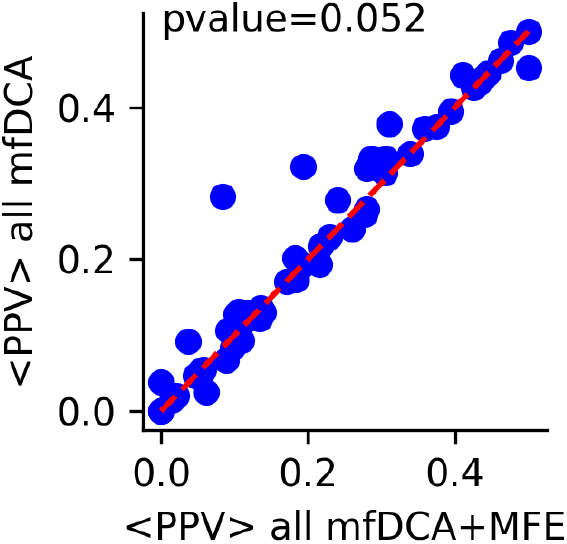
PPV comparison between biophysically informed and uninformed Mean Field DCA. The number of ranked coevolutionary scores used to compute PPV is set to half the RNA length.

**Fig. SI 4:**
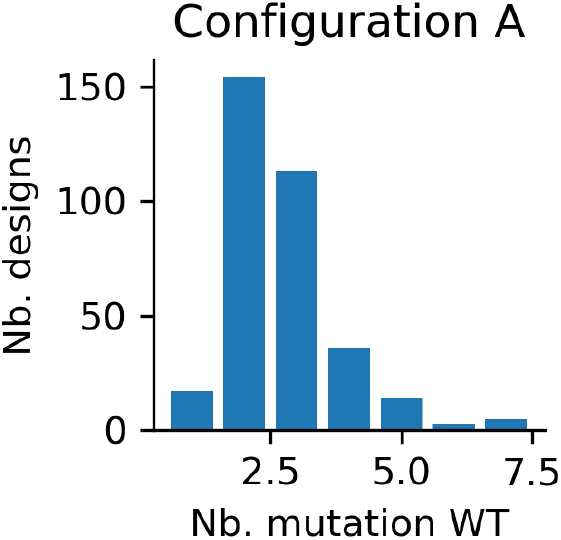
Artificial mutagenesis dataset for binding and folding. The barplot shows the number of selected random mutants of the artificial WT sequence designed for configuration A.

**Fig. SI 5:**
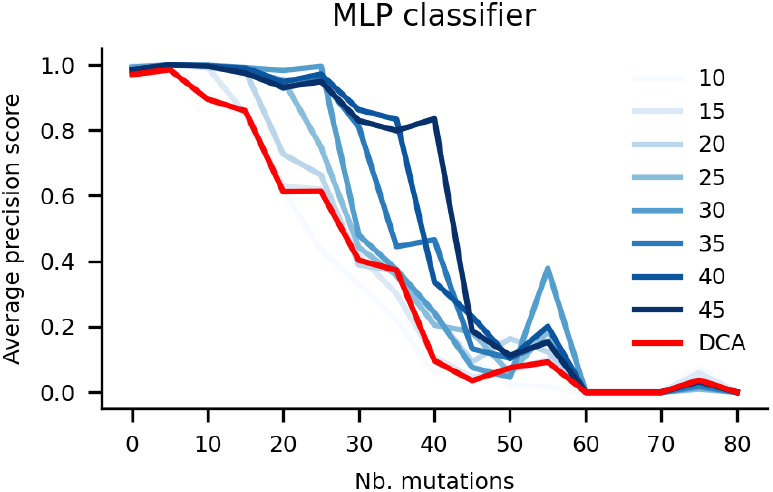
Average precision of MLP classifiers trained on increasing numbers of mutated sequences. Blue curves represent MLP models trained on 10–45 mutants (see legend). The red curve shows performance of a Potts model (DCA).

## Notes

### Competing Interest Statement

The authors have declared no competing interest.

### Summary of Updates

I have refine the model description.

